# PAWER: Protein Array Web ExploreR

**DOI:** 10.1101/692905

**Authors:** Dmytro Fishman, Ivan Kuzmin, Priit Adler, Jaak Vilo, Hedi Peterson

## Abstract

Protein microarray is a well-established approach for characterizing activity levels of thousands of proteins in a parallel manner. Analysis of protein microarray data is complex and time-consuming, while existing solutions are either outdated or challenging to use without programming skills. The typical data analysis pipeline consists of a data preprocessing step, followed by differential expression analysis, which is then put into context via functional enrichment. Normally, biologists would need to assemble their own workflow by combining a set of unrelated tools to analyze experimental data. Provided that most of these tools are developed independently by various bioinformatics groups, making them work together could be a real challenge. Here we present PAWER, the first online tool for protein microarray analysis. PAWER enables biologists to carry out all the necessary analysis steps in one go. PAWER provides access to state-of-the-art computational methods through a user-friendly interface, resulting in publication-ready illustrations. We also provide an R package for more advanced use cases, such as bespoke analysis workflows. PAWER is freely available at https://biit.cs.ut.ee/pawer.

## 1 Introduction

Protein microarray is the leading high-throughput method to study protein interactions [1], antibody specificity or autoimmunity [2]. In functional protein microarrays, full-length functional protein targets or protein domains are attached to the surface of the slide and then incubated with a biological sample that contains interacting molecules (e.g. autoantibodies) [3]. After molecules bind to their targets, labelling is done via secondary antibody with a fluorescent marker attached. Resulting fluorescent signal of high intensity indicates the reaction, which can be registered by the specialised scanner. The most popular microarray platforms (e.g. Human Proteome Microarray (HuProt), ProtoArray, NAPPA arrays, Human Protein Fragment arrays and Immunome arrays) allow to measure autoantibody reaction to thousands of unique human protein abundances simultaneously [4, 5].

Hundreds of studies that use different types of protein microarrays are conducted every year [6].All these studies largely depend on well executed data analysis. Usual analysis workflow starts with pre-processing of raw data obtained from GenePix Pro - one of *de facto* standard softwares used to read the microarrays [7]. The pre-processing step involves quality control and normalisation. It is followed by the differential protein analysis, in which protein reactivity levels that are significantly different between studied conditions are identified. These reactive protein levels are visualised, e.g. with boxplots. Finally, the results are interpreted using the body of prior knowledge via applying functional enrichment analysis tools. Setting up and executing these steps requires a lot of time and care from the researchers as each analysis step needs to be documented to ensure reproducibility.

Protein microarrays are similar to DNA microarrays as both technologies measure abundance of thousands of probes immobilised on the surface of the slide [8]. In the early days of protein microarrays research, this technological resemblance allowed practitioners to adapt methods and computational tools, originally developed for DNA microarrays [9]. However, a number of studies have shown that the same set of assumptions is not necessarily applicable to both types of microarrays, especially in terms of normalisation [5, 8, 9]. For example, in DNA microarrays the overall amount of signal is considered to be roughly the same between samples, while in protein microarrays only a small number of proteins are expected to show reactivity to probed serum. Applying quantile normalisation, that is usually utilised in DNA microarray analysis, may eliminate the relevant biological signal [8]. Thus, analytical pipelines tailored to protein microarrays are required in order to enable correct data analysis and consequently, biologically relevant results.

To date, three major tools for protein microarray analysis are Prospector, Protein Array Analyser (PAA) [10] and Protein Microarray Analyser (PMA) [5]. Prospector, provided by ThermoFisher Scientific, allows easy point and click analysis. However, it has not been updated since 2015, is a closed source software and runs only on the Windows 7 operating system [11]. PAA [10] builds on top of Prospector’s core functionality, and provides workflow customisation and tools for biomarker discovery in R. Although PAA is flexible and robust, it requires substantial programming skills from the user. PMA is a multi-platform desktop application, built in Java and published in 2018. It can be used via simple graphical user interface as well as executed from the command line. Although, PMA implements state-of-the-art normalisation and pre-processing strategies, working with it can be challenging, as to this date no relevant documentation is available. Moreover, neither of these protein microarray analysis tools allow the user to complete the full analysis workflow and thus demand external tools or packages, to be used downstream for visualisation and putting the results into context of previous knowledge.

Here, we present Protein Array Web Explorer (PAWER), the first web tool dedicated to analysing protein microarray data. PAWER incorporates the strengths of Prospector and PAA, while eliminating their major limitations. PAWER is suitable for experimental biologists who want to analyse their own data without the need to write code. PAWER has already been used for multiple projects, with the underlying R codebase central for analysis in two recent studies of APECED syndrome [2, 12].

## 2 MATERIALS AND METHODS

### 2.1 Key features

PAWER implements the following key features:

1. Public web service that can be used by anyone with protein microarray data in standard format
2. Interactive results table for convenient exploration of the results
3. Clear interactive visuals that can be downloaded in publication-ready formats
4. Parameterised algorithms at key steps (robust linear model (RLM), moderated T-test [13])
5. Downloadable intermediate results after each analysis step
6. Connection with g:Profiler [14] tool through its R package (gprofiler2) for fast enrichment analysis of differential protein features
7. An open source R package that the PAWER web service is built upon

### 2.2 Data upload and preprocessing

To start using PAWER, the user first needs to upload the fluorescent signal array readings - GenePix Results (GPR) files by either dragging and dropping files into the upload area or selecting them directly from the file system (via file upload window). Upon upload, PAWER automatically checks if submitted files come from the same platform and have the same extension. Detailed error message is shown in case any of these assumptions are not met. Once files have been successfully uploaded, the user is asked to select features that represent foreground and background intensities. In the case of ProtoArray and HuProt platforms, these values are chosen automatically, for other platforms user may have to manually search through the possible options from the drop-down menu. As soon as this is done, a global data matrix for the entire experiment is assembled from uploaded files using the limma R package [13]. Next, the background intensities are subtracted from the foreground values and signal from technical replicates is averaged. Resulting values are then log-transformed.

To reduce the technical noise in the data, we used a robust linear model trained on the set of protein features that are assumed to exhibit constant level of signal regardless of biological differences between samples. Such proteins are called positive controls and used in most of the platforms. Usually they are uniquely denoted in GPR files so that computer algorithms could identify them automatically. Hence, after files are uploaded, PAWER searches for such proteins and creates a list of potential positive controls. The list is then shown to the user for validation. User can alter it, by either removing or adding individual proteins. Robust linear model [8] is then used to predict the signal of control proteins based on their location (array and block) and type. In an ideal noise-free scenario, the resulting model will rely solely on protein type when predicting its signal, as any non-negative coefficient associated with array index or block id would indicate technical bias. In practice, unfortunately, noise is hard to avoid. Therefore, non-zero coefficients associated with individual protein arrays and blocks are subtracted from corresponding protein signals to remove technical bias. Data upload and normalisation steps normally take a few minutes, for example it took about 2 minutes to preprocess a dataset of size 770 Mb, with 100 samples.

After the normalisation step is complete, user can download the normalised data as a separate file. The file can be used as an input to other tools for additional analysis. Namely, in order to enable more elaborate cluster analysis, PAWER is linked to ClustVis [15]. ClustVis is a stand-alone online tool for cluster analysis and visualisation. ClustVis implements heatmaps and principal component analysis.

The final step in the PAWER data analysis pipeline is differential expression analysis which aims to identify proteins, which signal levels significantly differ between the sample groups. To execute this step, metadata (e.g. information about patients and controls) for each sample is required. User can either upload a separate metadata file or manually annotate every GPR file using the set of radio-buttons. The metadata file should contain only two columns: the list of filenames and corresponding sample groups.

### 2.3 PAWER output

Differential protein features are identified using a moderated t-test, implemented in the limma R package. To account for multiple testing, obtained p-values are adjusted by the Benjamini-Hochberg method. Proteins with adjusted p-values of less than 0.05 are considered significant and shown to the user in a table. User can filter the table by any value (e.g. protein name) and sort each field. By default it is sorted by the adjusted p-value. Results can be downloaded as a CSV text file for further analysis, as an Excel file to supplement a publication or as a PDF file to include into a presentation. To explore the underlying data distribution, individual protein expression values are visualised using interactive boxplots, which can be downloaded in a form of a publication-ready figure.

Additionally, enrichment of differential proteins is enabled by the gprofiler2 R package that provides interface for g:Profiler service [14]. g:Profiler gives functional enrichment results from a number of different categories, such as Gene Ontology [16], pathways and other structured data sources for instance KEGG [17], Reactome [18], Human Phenotype Ontology [19] and Human Protein Atlas [20]. The six most significant terms are visualised as a downloadable bar plot figure. The complete list of significantly enriched terms is accessible at the g:Profiler website.

### 2.4 Implementation

We developed the PAWER web service as a tool that covers all the necessary steps in protein microarray analysis. Its core has been implemented using R version 3.4.2, limma [21] (v. 3.34.4) for reading in the GPR format files and performing differential analysis, MASS [22] (v. 7.3.47), reshape2 [23] (v. 1.4.2) for normalisation and preprocessing of protoarray data and gprofiler2 [14] (v. 0.1.4) to enable protein identifier conversion and enrichment analysis. Web interface was implemented as a single page application using React.js and Redux architecture with node.js on the server side. Figures have been created and rendered with a help of D3.js [24] and DataTables libraries. Both the R package and the web server code are freely available under the GNU GPL v2. license.

### 2.5 Supported platforms

PAWER has been initially designed to support data produced mainly by ProtoArray and HuProt platforms. Later, support for the ArrayCam imaging system was added on request. Eventually, the decision was made to support as many platforms as possible by enabling customization at every step of the pipeline. Therefore, PAWER is in principle compatible with any protein microarray system or technology as long as the latter outputs text files with identical headers for each sample, and user knows several key properties of the system (background, foreground intensities and control proteins).

### 2.6 Comparison to existing tools

To the best of our knowledge there are four available tools, dedicated to protein microarray analysis – Prospector, PAA, PMA and PAWER. All the alternatives perform protein array specific normalisation and all but one (PMA) have capacity to identify potential biomarkers. The detailed comparison of the key features is highlighted in Table 1. Prospector was the first protein microarray analysis tool on the market, introduced by the Invitrogen company. It was originally developed for the Windows XP and later in 2015 updated to be compatible with Windows 7. Strict operating system dependency makes the number of potential Prospector users limited. In 2013 an R package, called PAA emerged [10]. Now users, independent from the platform, had an opportunity to design and apply custom analysis pipelines for their protein microarrays. At the same time, PAA requires users to be familiar with R programming language. Another tool, PMA - a Java desktop application, provided a graphical user interface and implemented cutting edge preprocessing techniques. However, it lacks documentation and does not allow for the integrated downstream analysis [5]. We developed PAWER as a freely accessible web service as an alternative way to analyse protein microarrays. PAWER has a user-friendly interactive graphical interface that helps researchers to apply standard protein microarray analysis pipeline with ease (Figure 1). Being comparable to PAA in its core strengths (protein specific normalisation and biomarker identification capabilities), PAWER uniquely helps to interpret the results of the analysis by providing detailed functional annotation of the identified differential proteins. Other key features are an interactive results table and accompanying attractive figures. Both figures and the table can be downloaded and used in the publications or scientific presentations. Notably, all the key steps of the analysis pipeline are well documented and presented on a separate help page.

**Table 1.** Comparison between currently available protein microarray analysis tools: Prospector, PAA, PMA and PAWER.

| Tool name | License | Last updated | Platform | GUI | Protein array specific normalisation | Potential Biomarker Identification | Detailed Functional Annotation | Downloadable visuals |
| --- | --- | --- | --- | --- | --- | --- | --- | --- |
| **Prospector** | No license specified | 2015 | Windows 7 Desktop application | + | + | + | — | — |
| **PAA** | BSD 3 | 2018 | R package | — | + | + | — | + |
| **PMA** | No license specified | 2018 | Java desktop application | + | + | — | — | — |
| **PAWER** | GNU GPL v2. | 2020 | Web server, R package | + | + | + | + | + |

**Figure 1.**
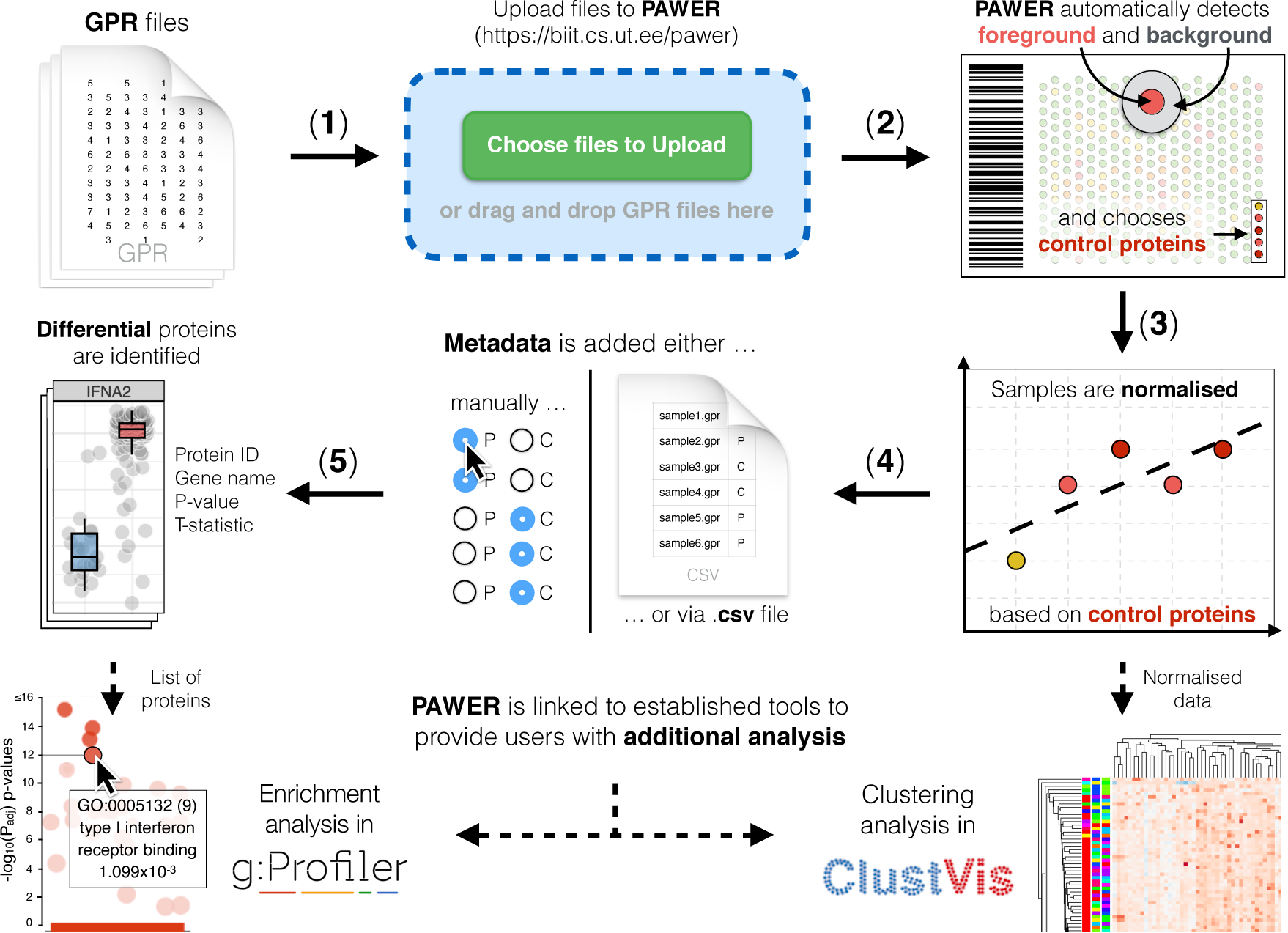
PAWER pipeline. Raw GPR files are uploaded to PAWER (**1**), then the system proceeds to identifying foreground and background intensities and a panel of control proteins that can be used for normalisation (**2**). Robust linear model is then used to estimate and remove the technical artifacts associated with each array and array block (**3**). Normalised data is then combined with sample metadata (**4**) to produce a list of differentially expressed proteins (**5**). PAWER is linked with two other tools (g:Profiler and ClustVis) to enable additional analysis, namely: protein enrichment analysis and cluster analysis of normalised expression values.

## 3 Conclusions

PAWER is the state-of-the-art protein microarray analysis pipeline with clean and intuitive web interface. The result of the analysis is presented as a searchable and filterable table. Interactive figures related to the table allow to explore reactivities in a more detailed manner. Both the table and the figures can be downloaded in various file formats, including in publication-ready visuals. In order to encourage further development of protein microarray analysis methods, both the R and the web application code are made openly available. PAWER has already been used in multiple projects, with the underlying R codebase central for analysis in two recent studies of APECED syndrome [2, 12].

To enable a closer interaction with our users and facilitate continuous improvement of PAWER, we have made available the PAWER feature roadmap, which can be accessed from the home page. It allows users to post feature requests and provide feedback.

## List of abbreviations

PAWER: Protein microarray web explorer
GPR: GenePix results
RLM: Robust linear model
PAA: Protein array analyser
PMA: Protein microarray analyser

## Data availability

PAWER web service is accessible online via https://biit.cs.ut.ee/pawer/. R package repository can be found via https://gl.cs.ut.ee/biit/paweR, while web interface repository is accessible at https://gl.cs.ut.ee/biit/pawer_web_client.

## Acknowledgements

The authors would like to acknowledge Pärt Peterson and Kai Kisand for introducing us to protein microarrays and their expert insight into the autoimmunity field. Also, we would like to thank Leopold Parts and Liis Kolberg for critical reading and comments on the manuscript. Special thanks go to Ms Boson for providing ideas that laid down the basis for the name and logo of PAWER.

## Author’s contributions

DF, PA analyzed protein microarray data and developed the R package. IK, PA, DF developed the web tool. DF wrote the paper with a contribution from JV, PA, HP and IK. HP supervised the project. All authors read, edited and approved the final manuscript.

